# Acellular starch domains in the endosperm of wheat and oat

**DOI:** 10.64898/2026.05.12.724639

**Authors:** Romuald Kosina, Paulina Tomaszewska, Łukasz Kochmański

**Affiliations:** Faculty of Biological Sciences, University of Wrocław, 51-148 Wrocław, Poland; Department of Genetics and Cell Physiology, Faculty of Biological Sciences, University of Wrocław, 50-328 Wrocław, Poland

**Keywords:** wheat, oat, development, microstructure, starch domains, endosperm cavity

## Abstract

The transformation of the free nuclear syncytium into cellular endosperm tissue with starch and protein accumulation is a well-established phenomenon, at least in the fruits of cereals of the Triticeae tribe. The present article demonstrates that there is considerable diversity inherent in this type of caryopsis morphogenesis. By examining various taxa (species, varieties, and cultivars) of wheat, oats, and some wild grasses, this research reveals significant deviations in endosperm morphogenesis from the typical state. A new developmental pattern of endosperm was identified, characterized by several distinctive features such as incomplete cellularization of the syncytium and starch accumulation within the acellular endosperm domains and the endosperm cavity. A large number of plastids were observed in the syncytium stage, which served as the basis for the later amyloplast stage. The acellular endosperm domains and the cavity domain exhibited connections at specific discontinuities in the modified aleurone layer surrounding the cavity. The peripheral parts of the caryopsis received fewer assimilates necessary for starch synthesis, which was attributed to their increased distance from the transfer system and a likely reduction in the efficiency of assimilate transport through the apoplast in these areas. The starch cavity volume constituted a few percent of the overall caryopsis volume, which could serve as a foundation for potential breeding improvements to enhance starch yields across different varieties.

## Introduction

The endosperm of cereals is the primary tissue involved in food production. The nutritional value of cereal products is mainly influenced by two final products of the assimilation process: protein and starch. To date, several concerns have been raised regarding the mechanisms underlying the development and morphogenesis of the endosperm of cereals belonging to various grass tribes. Although many studies have reported on the state of plastids in the syncytium, the alveolar stage of endosperm development, the cellularization pattern of its dorsal and ventral parts, the domain structure of the tissue, and the direction of assimilate transport (Mares et al., 1975; Simmonds and O’brien, 1981; Fineran et al., 1982; Batygina, 1987; Patrick and Offler, 2001; Sabelli and Larkins, 2009; Shewry et al., 2012; Olsen, 2020), our understanding of this phenomenon remains limited. Embryonic plastids occur primarily in functional megaspores and are involved in the formation of the *Polygonum*-type embryo sac in a single megaspore (Vijayaraghavan and Prabhakar, 1984). From this initial cell of the embryo sac, plastids divide and enter the egg cell, the central cell of syncytium, and the antipodals. These plastids contain starch (Kaltsikes et al., 1975; Batygina, 1987), which is activated at the early developmental stage to support the growing embryo sac. Embryonic plastids are the precursors to the starchy endosperm amyloplasts that develop later. Approximately 500 plastids/cell have been observed in wheat cv. Kolibri at about 4 days post anthesis (DPA) (Briarty et al., 1979). A question remains whether this initial pool of plastids is sufficient to fill the endosperm tissue and form mature grains. This depends on the efficiency of starch storage in amyloplasts and, therefore, ignoring the influence of the external environment, on the assimilative capacity of the organs proximal to the developing ovule, i.e., the flag leaf and all organs of the inflorescence and the pericarp tissue of caryopsis. The assimilative parenchyma of the pericarp, particularly abundant in the region of the vascular bundle, contains numerous starch grains that are utilized for endosperm formation (Morrison, 1976 Kosina, 2014). In cereals, endosperm development proceeds bidirectionally from the central cell walls in the form of alveoli and from the dorsal and ventral portions (Mares et al., 1975; Fineran et al., 1982). Although cytokinetic activity from the dorsal side appears to predominate over the ventral side, the tissue volume of both parts is also determined by cell growth, which is characterized by an elongated and intrusive pattern in the cylindrical endosperm (Tomaszewska and Kosina, 2018; Kosina and Tomaszewska, 2024).

In the endosperm tissue, a distinct line of contact-fusion is observed between the dorsal and ventral parts; however, in certain situations—hybrids and mutants—regions devoid of cells are noted (HÅKANSSON, 1953; Bosnes et al., 1987). It remains unclear whether this inhibition or lack of cell division occurs exclusively in such unusual states. As reported in the literature, the developmental principle follows a specific sequence: cellularization followed by starch accumulation in amyloplasts (Briarty et al., 1979; An et al., 2020). This raises the following three questions: (1) whether this developmental pattern is obligatory; (2) whether plastid and starch accumulation can occur in the coenocytic, acellular space; and (3) does grass endosperm morphogenesis differ from that commonly documented. Assessment of the potential for variation inherent in plants at different levels of organization (Johri, 1984; Briggs and Walters, 1986) indicates fluctuations in endosperm morphogenesis. For example, Tomaszewska and Kosina (2024) found that, at the syncytial level, the cytogenetic status of the oat embryo sac deviates from level 3*n* to 2*n* and can be interpreted as a *Polygonum*-type variant. Additionally, using common wheat accessions as an example, these authors demonstrated that the post-syncytial acellular space is not empty in the mature caryopsis, and that the free endosperm cavity domain participates in starch accumulation (Kosina and Tomaszewska, 2024). These observations indicate a specific potential for tissue variability at the developmental level. In this context, the present study used screening methods in accessions from different habitats to determine whether acellular morphogenesis occurs in other wheat and oat taxa, beyond hexaploid common wheat. Another important question is whether this altered morphogenesis pattern could play a critical role in cereal breeding.

## Materials and methods

Studies conducted on common wheat cultivars and hybrids grown under consistent soil and climatic conditions have shown that the frequency of acellular starch domains and the acellular endosperm cavity domain is specific to the studied accession(Kosina and Tomaszewska, 2024). This result established a fundamental framework for the screening analysis of accessions of various species, botanical varieties, and cultivated varieties of wheat and oat sourced from different locations and years of cultivation to investigate the presence of acellular endosperm and cavity domains. Table 1 provides a list of the analyzed accessions. Observations were conducted on cross-sections of the central part of the caryopsis, which was fixed in FAA fixative (formalin: acetic acid: ethanol) and cut with a microtome equipped with a freezing stage utilizing a TOS-11 selenium rectifier (V/O Medèksport, Moscow). In addition to assessing the total protein content, cross-sections of some accessions were stained with a 1% aqueous solution of bromophenol blue, washed in tap water, and mounted in glycerin on semi-permanent slides, in accordance with the method of Kaul et al. (1970). This method facilitated enhanced differentiation of the high-protein, starch-deficient subaleurone layer. The slides were photographed using an Amplival polarizing microscope (Carl Zeiss, Jena) as well as Olympus BX-50 and Zeiss Axio Imager M2 epifluorescence microscopes. An evaluation was conducted to assess the potential for increasing starch yield through its accumulation in the free space of the endosperm cavity for one caryopsis of *Triticum durum* cv. Candeal. This cultivar has medium-sized cavities compared to those previously examined in common wheat (Kosina and Tomaszewska, 2024). The volume ratio of the caryopsis and the endosperm cavity was measured on cross- and longitudinal sections prepared manually with a razor blade. The cavity shape was approximated to a cuboid, while the caryopsis was represented as a triaxial ellipsoid with volume V = 4/3Πa’b’c’ (where a’, b’, and c’ are the lengths of the semi-axes of the ellipsoid with axes a, b, and c, respectively).

**Table 1.** Accessions of cereals and wild grasses exemplified in the text.

| **Wheats** | **Donors** |
| --- | --- |
| *Triticum dicoccum* var. *rufum* | VIR |
| *Triticum durum* cv. Arancada | IGPB, Lublin, Poland |
| *Triticum durum* cv. Candeal | " |
| *Triticum aestivum* cv. Atlas 66 | Lincoln, University of Nebraska |
| *T. aestivum* cv. Lancota | " |
| *T. aestivum* cv. Purdue | " |
| *T. aestivum* cv. Octavia | Czechoslovakia |
| *Triticum fungicidum* | VIR |
| *Triticum kiharae* | VIR |
| Amphiploids and hybrid |  |
| *Triticum turanicum* var. *insigne*<br>x *Aegilops squarrosa* var. *stragulata* | Plant Germ-Plasm Institute<br>Kyoto |
| *Triticum dicoccum* cv. Vernal<br>x *Aegilops squarrosa* var. *stragulata* | " |
| *Triticum timopheevii*<br>x *Aegilops umbellulata* | " |
| *Triticum carthlicum* var. *darginicum*<br>x *T. durum* cv. Krasnodarskaja (F3) | R. Kosina |
| Oats |  |
| *Avena longiglumis* | BAZ |
| *Avena wiestii* | VIR |
| *Avena barbata* | BAZ |
| Amphiploids |  |
| *Avena abyssinica* x *A. strigosa* | NSGC, USA |
| *Avena barbata*<br>x *A. sativa* var. *nuda* | " |
| Wild grasses |  |
| *Bromus secalinus* | IPK (GRA 1040/81) |
| *Brachypodium distachyon* | IPK (GRA 788) |
| *Brachypodium silvaticum* | Wrocław, Poland (R. Kosina) |

NSGC - National Small Grains Collection, Aberdeen, Idaho, USA
VIR - Vavilov Institute of Plant Industry, St. Petersburg, Russia
BAZ - Bundesanstalt für Züchtungsforschung an Kulturpflanzen, Braunschweig, Germany
IPK – Institute of Plant Genetics and Crop Plant Research, Gatersleben, Germany
IGPB - Institute of Genetics and Plant Breeding, Agricultural Academy, Lublin, Poland

## Results and discussion

### Initial statements

The mother cell serves as a crucial unit in the development of a monosporic embryo sac. The functional megaspore provides the majority of plastids to the resultant daughter cells, including the egg cell, synergids, syncytium, and antipodals (Vijayaraghavan and Prabhakar, 1984). Male cells have a negligible contribution to plastid supply, for example, in rye, male cells provide approximately 2–12 plastids (Mogensen and Rusche, 2000). The plastids within the embryo sac can be classified as leucoplasts based on their content of starch grains, which can be observed under polarized light (Kaltsikes et al., 1975). This starch is utilized during the various developmental processes in the embryo sac, such as nuclear division, formation of alveoli, and cell division. In the syncytium, both in Triticeae amphiploid and oat, a substantial number of plastids are aggregated around the nuclei (Fig. 1a, b). In cereals from the Triticeae tribe, the presence of plastids, specifically leucoplasts, has been observed in the embryo sac in *Triticale*, in the egg cell, and in the zygote and syncytium. In the syncytium of the central cell, the number of starch grains and their size increase with the increasing number of nuclei (Kaltsikes et al., 1975). In wheat, leucoplasts were detected in the egg cell and syncytium (Batygina, 1987). Electron microscopy analysis revealed the presence of numerous plastids (2–3 μm diameter) with starch grains smaller than 0.1 μm diameter in the wheat syncytium at 2 DPA (Simmonds and O’BRIEN, 1981). In both *Triticale* and wheat, plastids accumulated around the nuclei. The a priori presence of plastids in the embryo sac syncytium is critical for their role in amyloplast morphogenesis in the developing endosperm of grasses. The mature endosperm tissue of cereals has two cell types: cylindrical and isolateral. The zones of cylindrical tissue, marked by oblique white lines in *Triticum durum* and the amphiploid *Avena* (Fig. 1c, d), exhibit considerable variation in size. Assimilates, primarily sucrose, are transported from the assimilating parts of the spikelets and pericarp through a transfer system, comprising the vascular bundle, pigment strand, nucellar projection, and endosperm cavity, directly to the central band of endosperm tissue situated along the caryopsis above the crease (FRAZIER and APPLANAIDU, 1965; Cochrane and Duffus, 1980; Patrick and Offler, 2001). This is highlighted in Fig. 1e and f and applies to cereals and other grasses with a similar caryopsis structure. Assimilates are transported symplastically through cells and apoplastically through cell walls and free spaces within the developing tissue. The endosperm cavity exemplifies one such apoplast structure. It may be inferred that the orderly, linear arrangement of walls in the cylindrical endosperm (Fig. 1c, d) facilitates such apoplastic transport, while the random, chaotic arrangement of walls in the endosperm on the lateral sides of the caryopsis slows it down. This arrangement of cells, i.e., cylindrical versus isolateral, in the starchy endosperm tissue results in high and low availability of assimilates to the central and peripheral parts, respectively. Large type A starch grains develop in the center of endosperm, while fewer and smaller starch grains develop in the distal parts. In these parts, the ratio between starch and protein also shifts in favor of protein. As shown in Fig. 1e-I, this phenomenon is observed in two varieties of common wheat and in *Bromus secalinus*, a grass that in the Middle Ages constituted a crop together with cultivated cereals. In wheat, the deficiency of assimilates in the distal parts is marked by a prominent protein-rich subaleurone layer, while in brome grass, this deficiency is characterized by the presence of distinctly smaller starch grains in the lateral parts of the caryopsis. Several studies have confirmed that the central transfer complex, including the cavity, plays a critical role in the transport of assimilates to the endosperm (Frazier and Appalanaidu, 1965; Cochrane and Duffus, 1980; Simmonds and O’BRIEN, 1981; Lim and Gifford, 1984; Patrick and Offler, 2001; CHATEIGNER-BOUTIN et al., 2021; Liu et al., 2022). Previous investigations have also emphasized the interaction of the symplast and apoplast in assimilate transfer. The efficiency of assimilate transport and starch accumulation depends on the vascular bundle, the most important element of the transfer complex. The development of a high-protein subaleurone layer with deficiency in starch grains correlates with the bundle, which has fewer vascular cells (Kosina, 1988). A similar development is observed in the distal, low-starch grains in the spikelets and spike (Nicolas et al., 1985; Stoddard, 1999).

**Fig. 1.**
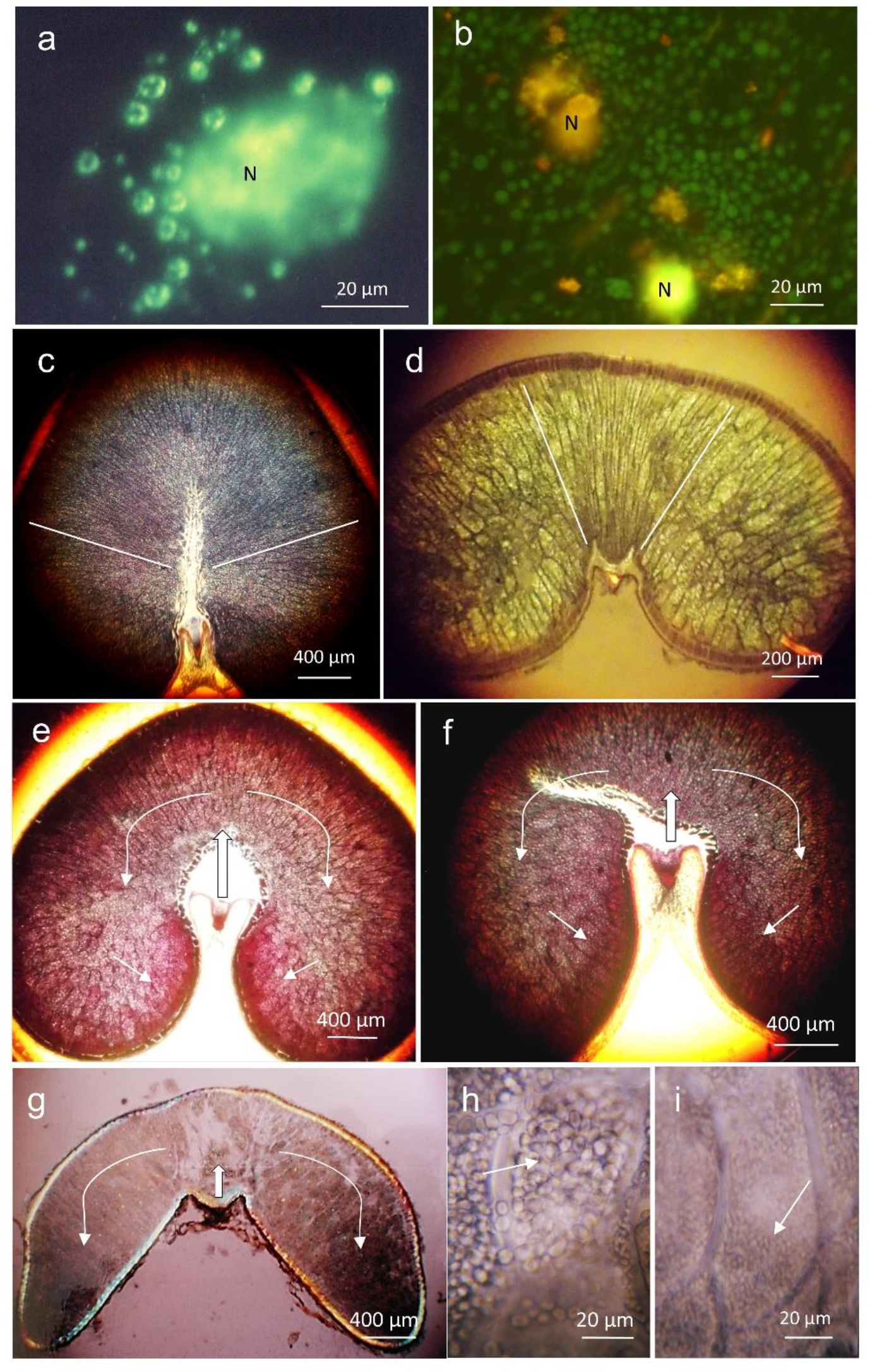
Young syncytial plastids (**a, b**) and assimilate transport routes (**c**-**i**). **a** – **a** – *Triticum turanicum* var. *insigne* x *Aegilops squarrosa* var. *strangulata* (fluorescein isothiocyanate fluorescence; N – nucleus, plastids clustered around the nucleus); **b** – *Avena barbata* (acridine orange fluorescence, N: nucleus); **c** – *Triticum durum* cv. Arancada (white lines border a broad domain of cylindric dorsal endosperm); **d** – *Avena abyssinica* × *A. strigosa* (white lines border a narrow domain of cylindric dorsal endosperm); **e** – *Triticum aestivum* cv. Atlas 66 (the main direction of assimilate transport – block arrow, secondary transport – curved arrows, high-protein subaleurone layer below the cavity – small arrows); **f** – *T. aestivum* cv. Purdue (marked as in **e**); **g, h, i** – *Bromus secalinus*; **g** (marked as in **e**); **h** and **i** – arrows show large amyloplasts above the transfer system (**h**) and small amyloplasts in the distant lateral parts (**i**). **c, e, f** – stained with bromophenol blue; **c** – polarized light (partly crossed nicols).

### Dorsal-ventral endosperm contacts and acellular domains in the starchy endosperm

In cereals, the endosperm cell structures develop at the embryo sac wall as alveoli containing cytoplasm and nuclei (Mares et al., 1975). A cross-sectional analysis of a young caryopsis shows the development of dorsal and ventral tissues, which culminates in the formation of alveoli toward the center of the central cell of the embryo sac. The contact of alveoli of both parts leads to the formation of a double cell wall. This further reveals the line of contact between the dorsal and ventral parts (Fineran et al., 1982). However, these authors consider the process of cell contact formation to be an open question. Examples of such contacts are shown in Fig. 2a, b, which illustrate equivalence in tissue volume on the dorsal and ventral sides, both in the central and lateral parts of the caryopsis. Bosnes et al. (1992) described a similar situation for barley *sex* mutants. However, the free space in the center of the central cell is not always closed. This phenomenon has been documented in studies on barley hybrids (HÅKANSSON, 1953) and the hexaploid *Triticale* (Pena et al., 1982). Fig. 2c shows the free space between endosperm cells. The cross-section was obtained from a caryopsis of the synthetic amphiploid *Triticum dicoccum* × *Aegilops squarrosa*, which was developed by M. Tanaka (Kyoto) in 1959. This amphiploid was housed in the collection of the Plant Germ-Plasm Institute in Kyoto. Following the acquisition of seed material from this institution, the amphiploid was cultivated for many years in the grass collection maintained by R. Kosina. Therefore, it can be assumed that the amphiploid exhibits a degree of developmental stability resulting from natural selection during prolonged cultivation. Because the starchy endosperm cells in contact with the free space are not compressed by other cells, their free-growing walls are rounded. Dorsal-ventral contacts are distinct (Fig. 2a, b) due to the contact of two developed walls; however, there is no double wall formation between alveoli during cytokinesis. Free walls are characterized by the specificity of their outer layers in plant organs; when these layers come into contact or merge, they are altered due to an increase in the content of callose and unesterified pectins, for example, *adherent1* mutation in maize (Sinha and Lynch, 1998). It is also plausible that the distinct contacts observed in the endosperm may have a similar underlying mechanism.

**Fig. 2.**
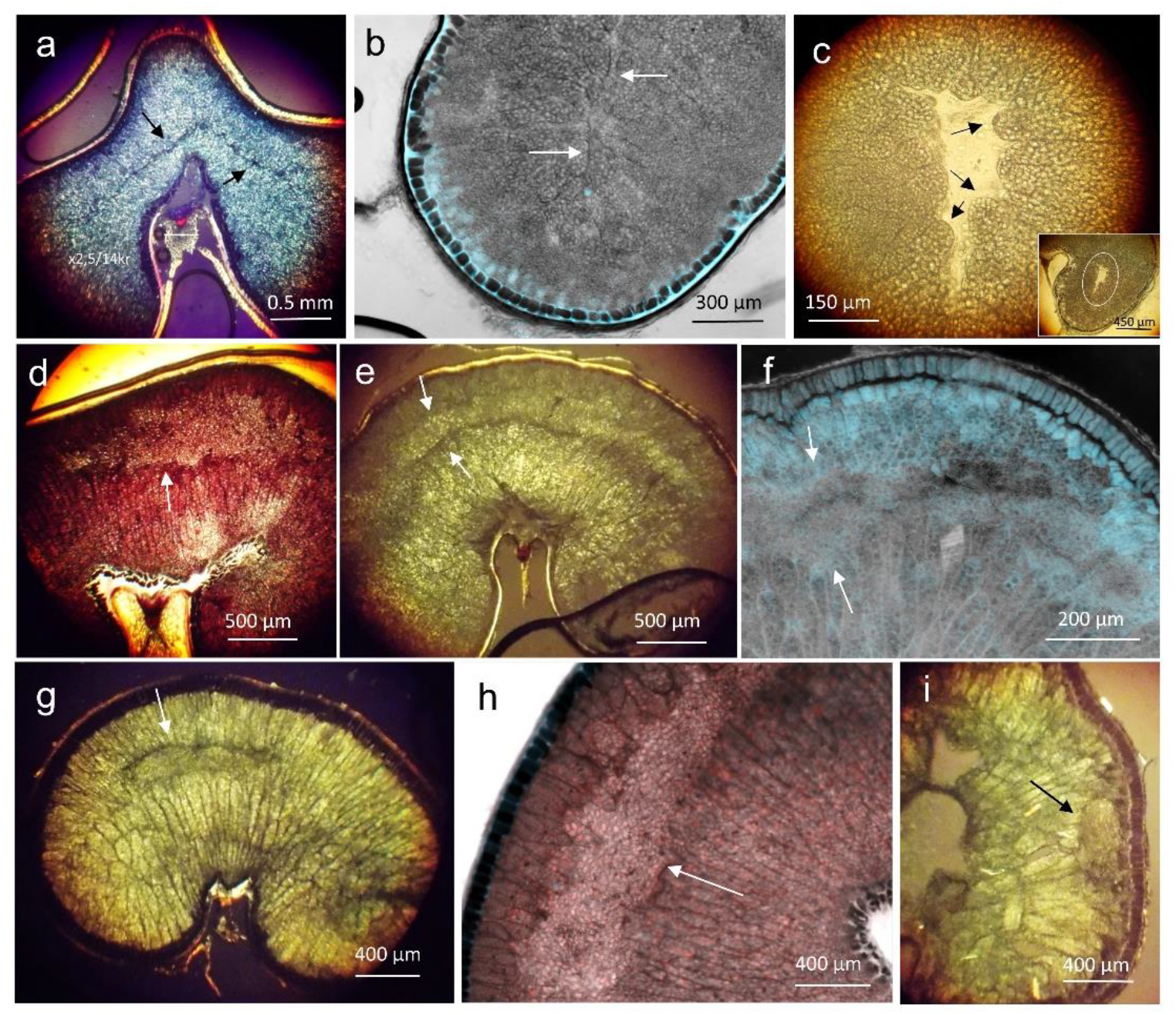
Area of cellular contact of dorsal and ventral endosperm (**a**-**c**) and acellular dorsal and lateral starchy domains (**d**-**i**). **a** – *Triticum carthlicum* var. *darginicum* × *T. durum* cv. Krasnodarskaja (F3 generation, arrows indicate the dorsal-ventral contact in the central part of the caryopsis); **b** – *Triticum dicoccum* cv. Vernal × *Aegilops squarrosa* var. *strangulata* (contact of endosperm cells in the lateral part of the caryopsis, at the crease); **c** – *T. dicoccum* cv. Vernal × *Ae. squarrosa* var. *strangulata* (empty space in the endosperm surrounded by round-walled cells; located in the lateral part of the caryopsis – see inset); **d** – *T. aestivum* cv. *Purdue* (arrow indicates the acellular dorsal domain); **e** – *Triticum fungicidum* (extensive, arcuate acellular dorsal domain between arrows); **f** – *Avena longiglumis* (dorsal domain between arrows), **g** – *Avena abyssinica* × *A. strigosa* (lenticular dorsal domain); **h** – *T. aestivum* cv. Lancota (lateral domain); **i** – *Avena barbata* × *A. sativa* var. *nuda* (lenticular lateral domain under the subaleurone layer). **a, e, g, i** – partially polarized light; **b, f** – autofluorescence; **d, h** – stained with bromophenol blue.

Starch can accumulate in amyloplasts in the free spaces of the endosperm, and this developmental event was first observed by Kosina and Tomaszewska (2024) in some common wheat cultivars. Such acellular domains are shown in Fig. 2d-i. Examples are provided for some wheat species and cultivars and oats. Cross-sections of the caryopsis reveal the domains are located in the dorsal part opposite the transfer system and in the lateral parts. An asymmetry is observed in the volume of tissue located beyond the domains toward the periphery of the caryopsis and the part directed toward the transfer system and the cavity. The latter is thicker, with a larger number of cells. Thus, a discrepancy is noted between the location of the interface (Fig. 2a) and the location of the domain (Fig. 2d). This condition suggests the likelihood of local incompletion or delay of cytokinesis in the peripheral parts of the caryopsis.

### Variability of endosperm cavity volume and discontinuity of the modified aleurone layer (MAL)

In the grains of grasses of the Triticeae tribe and other tribes with grains possessing a similar tissue pattern, the endosperm cavity is the sole region in direct contact with the starchy endosperm. As mentioned earlier, this cavity serves as a component of the transfer system for assimilates and does not function as a starch accumulation site. Kosina and Tomaszewska (2024) found that starch accumulation in the cavity occurs with significantly higher frequency in some common wheat cultivars. In mature caryopses, the observed cavities vary in volume, from negligible to large. A pertinent example exhibiting this variation is shown for two cultivars of *T. aestivum*, namely Octavia and Lancota (Fig. 3a, b). This finding demonstrates that the large cavity domain has considerable potential for starch accumulation. Kosina (1984) reported significant variability in cavity size among species such as *T. monococcum* and *T. durum*. This observation was further confirmed by studies conducted by Kosina and Tomaszewska (2024) for other common wheat cultivars and hybrids.

**Fig. 3.**
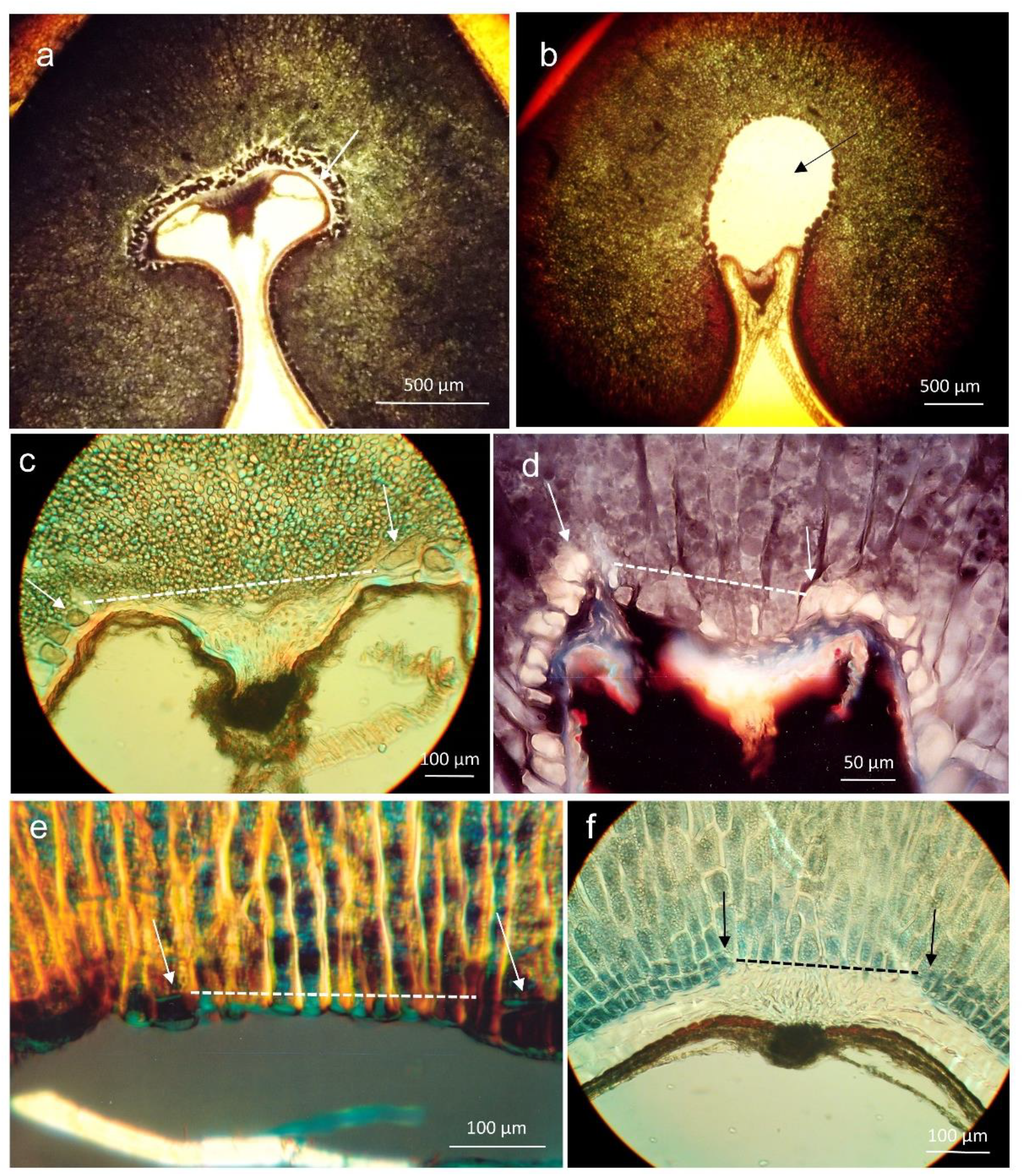
Variability of the size of endosperm cavities (**a, b**) and discontinuities of the modified aleurone layer (MAL) (**c**-**f**). **a** – *Triticum aestivum* cv. Octavia (a very narrow, small cavity – arrow); **b** – *T. aestivum* cv. Lancota (a large, voluminous cavity – arrow); **c** – *T. timopheevii* × *Aegilops umbellulata* (MAL discontinuity – a broken line, and aleurone cells at the edges of the gap – arrows); **d** – *Avena wiestii*, **e** – *Brachypodium silvaticum*; **f** – *B. distachyon*; **d, e, f** – designation as in **c. a, b, f** – stained with bromophenol blue; **c, e** – partially polarized light; **d** – autofluorescence.

The aleurone layer forms a boundary between the cavity and the starchy endosperm and is different from the layer surrounding the endosperm beneath a seed coat. It is often multi-layered and comprises irregularly shaped cells; these cells exhibit intrusive growth towards the dorsal layer. Discontinuities in this layer creating direct contacts between the endosperm cavity contents and the starchy cells are a rare feature (Tomaszewska and Kosina, 2018; Kosina and Tomaszewska, 2024).

The aleurone layer of the cavity is described as the MAL (EVERS and MILLAR, 2001). Here, a critical aspect is the formation of discontinuities within this layer. This developmental anomaly has been documented for several grasses, including wheat and oats (Fig. 3c-f). The degree of contact between the cavity and starchy tissue exhibits variability, with such variations most frequently observed in the central part, toward the dorsal endosperm. The endosperm cavity stretches along the caryopsis, above the nucellar attachment. It extends from the position of the coleorhiza around the radicle to the end of the crease. Consequently, it adopts a sac-like configuration, which may be perforated by MAL discontinuities throughout its entire length. This MAL structure may play a significant role in the bilateral exchange between the cavity and the endosperm during the phase of incomplete cytokinesis as well as during ongoing starch synthesis and accumulation.

### Acellular starch domain in the cavity and callus-like cells bordering acellular domains

MAL discontinuities in contact with the acellular space in the starchy endosperm tissue allow syncytial amyloplasts to enter the cavity (Fig. 4a, b), leading to complete (Fig. 4c-e) or partial (Fig. 4f) filling of the cavity. Occasionally, a directional arrangement of amyloplasts is observed (Fig. 4b). A noteworthy aspect is that the accumulation of amyloplasts and the subsequent deposition of starch in them can occur not only in well-developed, large caryopses but also in small, thin grains, as observed in *T. dicoccum* var. *rufum* (Fig. 4e and inset). Because plastid and starch accumulation occurs in the free space, endosperm cells in contact with this free space form oval walls, as noted in callus tissue in wound areas or in *in vitro* cultures. Fig. 4g-h shows such walls for *T. kiharae* and *T. aestivum* cv. Purdue. “Callus-like” endosperm cell development has also been demonstrated in barley mutants (BOSNES et al., 1997, 1992) and in *in vitro* cultures of *Zea* (Olsen, 2020). Starch cavity filling has been observed in T. *dicoccum* var. *chwamlicum* with a frequency of up to 30% (Kosina, 1984); and *T. aestivum* cultivars, namely Nap Hal, Lancota, and Purdue, with a frequency of 1%, 5%, and 14%, respectively (Kosina and Tomaszewska, 2024).

**Fig. 4.**
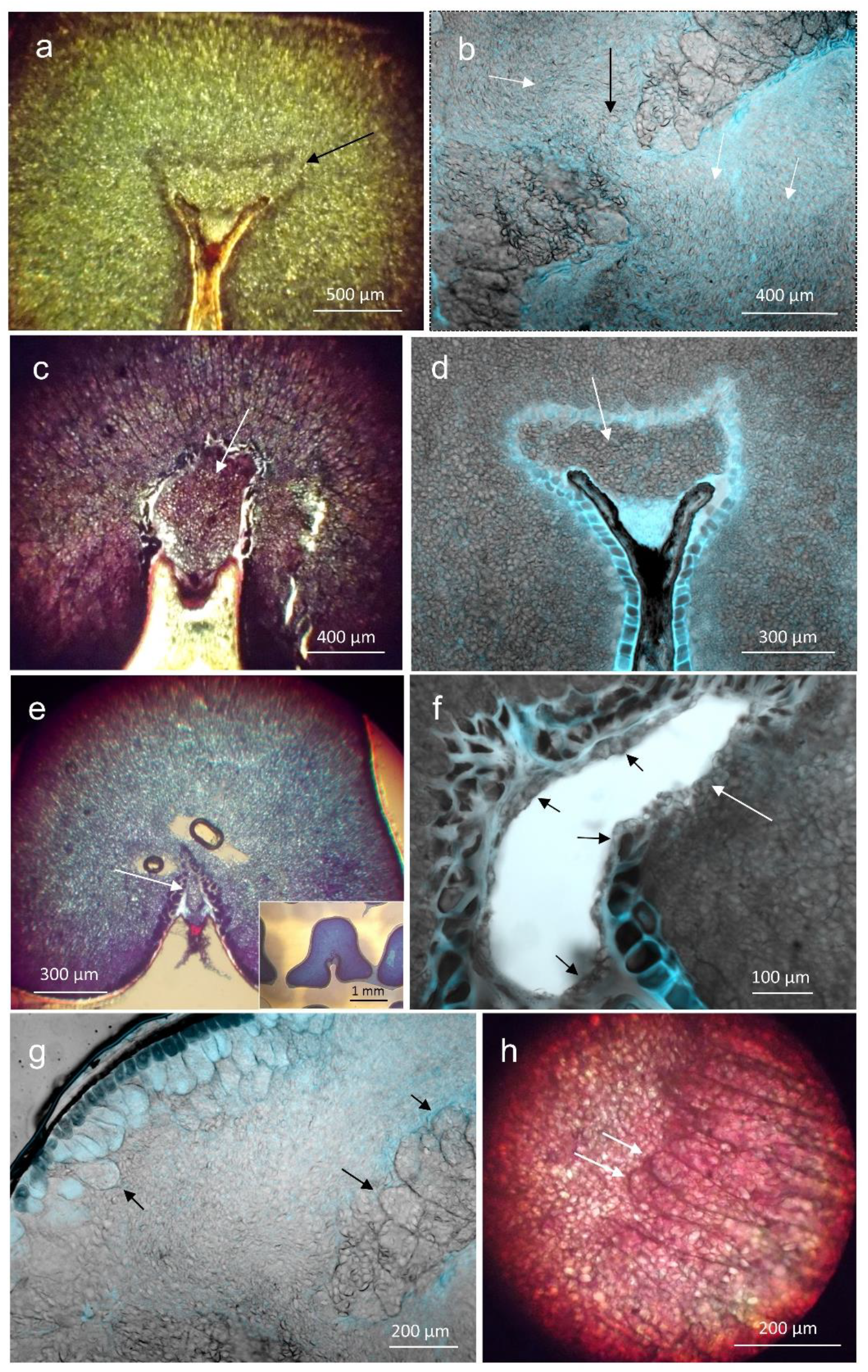
Contacts of acellular cavity domains with the cellular endosperm (**a**) or its acellular domain (**b**), starch accumulation in the cavity (**c**-**f**), and callus-like cells at the border of endosperm acellular domains (**g**-**h**). **a** – *Triticum dicoccum* cv. Vernal × *Aegilops squarrosa* (connection of cavity starch with endosperm tissue outside the cavity – arrow); **b** – *T. kiharae* (connection of acellular domains of cavity and endosperm – black arrow, directional arrangement of starch grains – white arrows); **c** – *T. aestivum* cv. Purdue (cavity filled with starch); **d** – *T. dicoccum* cv. Vernal × *Ae. squarrosa* (cavity filled with starch); **e** – *T. dicoccum* var. *rufum* (small cavity in the lean caryopsis filled with starch, inset showing a cross-section of the caryopsis typical for the tested sample); **f** – *T. aestivum* cv. Purdue (cavity with few starch grains adjacent to the MAL – black arrows and a gap in the MAL – black arrow); **g** – *T. kiharae* (callus-like cells bordering the acellular domain – arrows); **h** – *T. aestivum* cv. Purdue (callus-like ends of starchy cells bordering the acellular domain – arrows). **a** – polarized light; **b, d, f, g** – autofluorescence; **c, e, h** – stained with bromophenol blue.

### Comparison of cavity and caryopsis volumes

It is essential to evaluate the variations in endosperm morphogenesis discussed here, particularly the accumulation of amyloplasts and starch in the additional domain of the cavity, in terms of their practical significance and relevance to cereal breeding practices. A pertinent concern in this context is the extent to which the free space of the cavity increases starch yield per hectare. As shown in Fig. 5a, starch volume in the acellular domains within the endosperm tissue is significantly greater than cavity volume. In these domains, assimilates are used for starch synthesis rather than for cytokinesis. The cavity is the only “free cell to occupy.” An attempt was made to estimate the ratio of the caryopsis volume (endosperm with the fruit-seed coat) to the volume of the cavity by using the example of a single caryopsis of T. *durum* cv. Candeal, a cultivar of Argentine origin. The analyzed caryopsis has a medium-sized cavity compared to those in wheat (Kosina, 1984; Kosina and Tomaszewska, 2024). The caryopsis shape was approximated to a three-axis ellipsoid, while the cavity shape was described as a cuboid, without considering the shape and volume of the cavity located beneath the embryo. Based on this approximation and calculation, the cavity constitutes approximately 3.8% of the volume of the entire caryopsis.

**Fig. 5.**
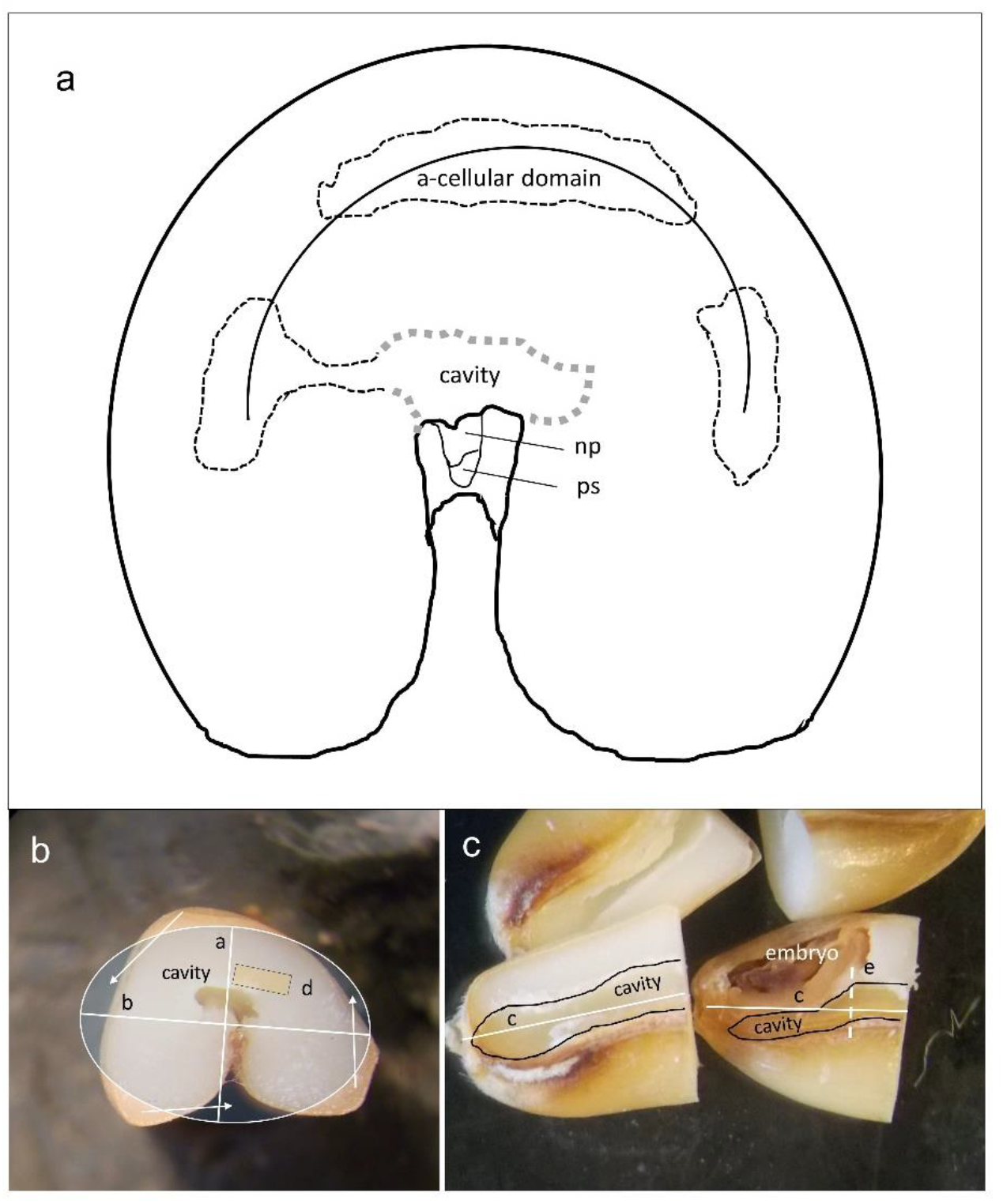
Diagram of the distribution of acellular starch domains in a cross-section of a caryopsis (**a**) and analysis of the caryopsis and cavity volume in *Triticum durum* cv. Candeal (**b**-**c**). **b** – cross-section of a caryopsis (a, b – short axes of the ellipsoid; d – rectangle approximating the transverse shape of the cavity, auxiliary arrows – approximation of the caryopsis shape to an ellipsoid); **c** – longitudinal section of a caryopsis – two parts after cutting in the **b** plane (c – long axis of the ellipsoid, e – dashed line outside the embryo as the starting point for measuring the cavity length, black line indicates the cavity shape on the longitudinal section); **a**: ps: pigment strand, np: nucellar projection.

In lean, sparsely filled caryopses (Fig. 4e), the presence of amyloplasts throughout the entire volume of the small cavity suggests that this morphogenesis is independent of yield-determining environmental factors. Cultivars exhibiting cavities that are almost closed by the MAL layer and adjacent nucellar projection (Fig. 3a) are unlikely to be used as viable candidates for breeding programs aimed at enhancing starch yield per caryopsis.

It can be assumed that, compared to the calculated value of 3.8%, the cavity proportion in many breeding candidates might be actually higher. Currently, the extent to which the reported novel type of endosperm morphogenesis could be considered an environmentally modified event remains unclear. The observation that starch cavities occur with greater frequency in some cases (Kosina, 1984; Kosina and Tomaszewska, 2024) indicates the possibility of achieving selective progress.

The analyzed accessions reveal a new type of endosperm morphogenesis that deviates from the previously established forms. This unique process is characterized by two salient features: incomplete cellularization of the central cell syncytium and an accumulation of excess starch in the cavity. Incomplete syncytium cellularization was observed in rice due to mutation (Tonosaki et al., 2021). The results noted for free spaces in the endosperm of barley hybrids (HÅKANSSON, 1953) and *Triticale* (Pena et al., 1982) support the possibility of impaired cellularization; however, in these examples, the free spaces are associated with unstable forms (mutation, hybridity, or unstable alloploid). Although the materials analyzed in the present study also include some unstable forms, e.g., wheat octoploids (*T. kiharae*), alloploids, and an F3 hybrid, several accessions are stable types, species, varieties, and cultivars. Based on our findings, the previously reported developmental sequence of cereal endosperm, i.e., cellularization → starch accumulation, should be expressed as incomplete cellularization → starch accumulation. This developmental pathway provides the plant with additional energy resources that can be used for starch synthesis and accumulation, instead of utilization for cytokinesis and wall growth. Starch synthesis is likely initiated in plastids present at the syncytium stage, or starch accumulation might persist for a longer duration. Both endosperm cellularization and plastid division and starch synthesis within them require substantial energy input; this energy is supplied through the process of programmed cell death (PCD) in pericarp tissues (Sabelli and Larkins, 2009; Dominguez and Cejudo, 2014). PCD in the pericarp and endosperm is correlated with starch accumulation in the endosperm (Li et al., 2010). This phenomenon was initially confirmed by Morrison (1976) based on the starch pool in the mother tissue of the wheat caryopsis; the cross-cell pericarp layer contains many starch grains from approximately 10 DPA to 17 DPA, which disappear only around 21 DPA. Thus, whether starch accumulation in the acellular domains of the endosperm and cavities is influenced by the rate and duration of accumulation is a possibility that requires further validation. This possibility is indicated by genotypic variability in these parameters between accessions. For example, an analysis of the starch accumulation rate and the caryopsis filling rate in two wheat cultivars with varying starch content, Annong9912 (54%) and E28 (69%), showed that the E28 cultivar had a higher amylopectin and amylose content due to earlier initiation of starch accumulation coupled with a higher rate of accumulation (Wang et al., 2014). Li et al. (2023) showed that the varietal differences between Xindong 18 and Xindong 22 in 1000-grain weight and in sucrose content and sucrose synthase activity were also maintained under drought stress conditions, with the latter variety exhibiting a higher 1000-grain weight.

Assimilate flow and starch accumulation depend on two compartments in the tissue: the symplast and apoplast. Although the apoplast volume in the wheat grain constitutes merely a few percent, sucrose concentration in the endosperm cavity, an important component of the apoplast, was found to determine the growth rate of the caryopsis (Lim and Gifford, 1984; Patrick and Offler, 2001). The translocation of assimilates into the cavity occurs via the parent tissues through the transfer system, i.e., the vascular bundle, pigment strand (ps), and nucellar projection (np). Two distinct morphotypes of np tissues have been identified, and their architecture can accelerate or delay the flow of assimilates (Kosina, 2014). With extended accumulation of starch, the sole remaining free compartment of the seed, i.e., the endosperm cavity, becomes the primary site for starch storage. The cavity contains arabinoxylans and hydroxycinnamic acids from the epidermis of the nucellus, in the form of a hydrated gel (CHATEIGNER-BOUTIN et al., 2021), which probably enables starch accumulation in it at the last stage of caryopsis filling.

### Concluding remarks

The following conclusions can be drawn from observations of transverse sections of the caryopsis. Numerous plastids, specifically leucoplasts, present in the cells of the embryo sac, including the central cell syncytium, serve as the foundation for the morphogenesis of amyloplast in the endosperm cells. Assimilates for starch synthesis in amyloplasts and proteins are transported from the transfer complex (vascular bundle, ps, and np) through the hydrophilic domain of the endosperm cavity. The architecture of the endosperm tissue facilitates a more efficient transport of assimilates to the center of the caryopsis due to its cylindrical cell structure rather than to the lateral parts comprising random arrangements of the apoplastic cell walls. This phenomenon is supported by the observation that the region of endosperm tissue located adjacent to the crease in the lateral parts of the caryopsis has the thickest, protein-rich, starch-deficient subaleurone layer. Throughout the growth of the endosperm on both dorsal and ventral sides, the interface between the cells of both parts is maintained, with less frequent occurrence of free space between them. The discovery of acellular domains in the central and lateral regions indicates that the cellularization process in the central cell is inhibited at an earlier stage, leading to the accumulation of plastids and starch in the coenocytic space. The MAL surrounding the cavity exhibits discontinuity, likely facilitating the transport of assimilates to the central cell and acellular domains. In the absence of mechanical pressure from the surrounding tissue, starchy endosperm cells in contact with the acellular domains exhibit wall development typical of callus tissue. During the final stage of accumulation of assimilates in the caryopsis, a specific proportion of amyloplasts containing starch accumulates in the cavity domain. This process is facilitated by the hydrophilic state of the cavity. The novel type of endosperm morphogenesis described above is a characteristic of some wheat and oat accessions, which creates new opportunities for improving starch yield during breeding. Starch storage in the free cavity domain can increase starch yield per caryopsis by several percent.

## CRediT authorship contribution statement

**RK:** scientific conception, investigation, data curation, manuscript writing. **PT:** fluorescence microscopy data analysis and corresponding figure preparation, manuscript review and enhancement. **ŁK:** collection of data on *Bromus secalinus*. All authors have read and approved the final version of the manuscript.

## References

An, L., Tao, Y., Chen, H., He, M., Xiao, F., Li, G., Ding Y., Liu, Z., 2020: Embryo-endosperm interaction and its agronomic relevance to rice quality. Front. Plant Sci. 11, 587641. DOI: 10.3389/fpls.2020.587641.

Batygina, T.B., 1987: Chlebnoe zerno. Atlas. Nauka, Leningrad.

Bosnes, M., Harris, E., Aigeltinger, L., Olsen, O.-A., 1987: Morphology and ultrastructure of 11 barley shrunken endosperm mutants. Theor. Appl. Genet. 74, 177–187.

Bosnes, M., Weideman, F., Olsen, O.-A., 1992: Endosperm differentiation in barley wild-type and sex mutants. Plant J. 2, 661–674. DOI: 10.1111/j.1365-313x.1992.tb00135.x.

Briarty, L. G., Hughes, C.E., Evers, A.D., 1979: The developing endosperm of wheat — a stereological analysis, Ann. Bot. 44, 641–658. DOI: 10.1093/oxfordjournals.aob.a085779.

Briggs, D., Walters, S.M., 1986: Plant variation and evolution. Cambridge University Press, Cambridge.

Chateigner-Boutin, A.-L., Alvarado, C., Devaux, M.-F., Durand, S., Foucat, L., Geairon, A., Grélard, F., Jamme, F., Rogniaux, H., Saulnier, L., Guillon, F., 2021: The endosperm cavity of wheat grains contains a highly hydrated gel of arabinoxylan. Plant Sci. 306, 110845. DOI: 10.1016/j.plantsci.2021.110845.

Cochrane, M.P., Duffus, C.M., 1980: The nucellar projection and modified aleurone in the crease region of developing caryopses of barley (Hordeum vulgare L. var.distichum). Protoplasma 103, 361–375. DOI: 10.1007/BF01276962.

Dominguez, F., Cejudo, F.J., 2014: Programmed cell death (PCD): an essential process of cereal seed development and germination. Front. Plant Sci. 28. DOI: 10.3389/fpls.2014.003.

Evers, T., Millar, S., 2002: Cereal grain structure and development: Some implications for quality. J. Cereal Sci. 36, 261–284.

Fineran, B.A., Wild, D.J.C., Ingerfeld. M., 1982: Initial wall formation in the endosperm of wheat, Triticum aestivum: a reevalution. Can. J. Bot. 60, 1776–1795.

Frazier JC, Appalanaidu B (1965: The wheat grain during development with reference to nature, location, and role of its translocatory tissues. Am. J. Bot. 52, 193–198.

Håkansson, A., 1953: Endosperm formation after 2x, 4x crosses in certain cereals, especially in Hordeum vulgare. Hereditas 39, 57–64.

Johri, B.M., 1984: Embryology of Angiosperms. Springer, Berlin.

Kaltsikes, P.J., Roupakias, D.G., Thomas, J.B., 1975: Endosperm abnormalities in Triticum – Secale combinations. I. x Triticosecale and its parental species. Can. J. Bot. 53, 2050–2067.

Kaul, A.K., Dhar, R.D., Swaminathan, M.S., 1970: Microscopic and other dye-binding techniques of screening for proteins in cereals. In: Improving Plant Protein by Nuclear Techniques, Proc. Symp. Vienna, I.E. & FAO, pp 253–262.

Kosina, R., 1984: Morphology of the crease of wheat caryopsis and its usability for identification of some species – a numerical approach. In: van Zeist W, Casparie WA (eds) Plants and Ancient Man. A.A. Balkema, Rotterdam, pp 177–191.

Kosina, R., 1988: Relationship between xylem bundle and subaleurone endosperm layer in wheat tetraploids caryopses. Hod. Rośl. Aklim. Nasienn. 32, 235–237.

Kosina, R., 2014: On caryopsis xylem and its interactions. In: Berhardt LV (ed) Advances in Medicine and Biology 75. Nova Science Publishers, New York, pp 43–63.

Kosina, R., Tomaszewska, P., 2024: Multi-character assessment of caryopsis in diverse common wheat germplasm, including high protein breeds. bioXarchiv. 10.1101/2024.10.26.620432.

Li, C.Y., Li, W.H., Li, C., Gaudet, D.A., Laroche, A., Cao, L.P., Lu, Z.X., 2010: Starch synthesis and programmed cell death during endosperm development in triticale (× Triticosecale Wittmack). J. Integr. Plant Biol. 52, 602–615.

Li, C., Fu, K., Guo, W., Zhang, X., Li, C., Li, C., 2023: Starch and sugar metabolism response to post-anthesis drought stress during critical periods of elite wheat (Triticum aestivum L.) endosperm development. J. Plant Growth Regul. DOI: 10.1007/s00344-023-10930-3.

Lim, C.H., Gifford, R.M., 1984: Accumulation and conversion of sugars by developing wheat grains: V. The endosperm apoplast and apoplastic transport. J. Exp. Bot. 35, 58–73. DOI: 10.1093/jxb/35.1.58.

Liu, J., Wu, M.-W., Liu, C.-M., 2022: Cereal Endosperms: Development and Storage Product Accumulation. Annu. Rev. Plant Biol. 73, 255–291.

Mares, D.J., Norstog, K., Stone, B.A., 1975: Early stages in the development of wheat endosperm. I. The change from free nuclear to cellular endosperm. Aust. J. Bot. 23, 311–326.

Mogensen, H.L., Rusche, M.L., 2000: Occurrence of plastids in rye (Poaceae) sperm cells. Am. J. Bot. 87, 1189–1192.

Morrison, I.N., 1976: The structure of the chlorophyll-containing cross cells and tube cells of the inner pericarp of wheat during grain development. Bot. Gaz. 137, 85–93.

Nicolas, M.E., Gleadow, R.M., Dalling, M.J., 1985: Effect of post-anthesis drought on cell division and starch accumulation in developing wheat grains. Ann. Bot. 55, 433–444. DOI: 10.1093/oxfordjournals.aob.a086922.

Patrick, J.W., Offler, C.E., 2001: Compartmentation of transport and transfer events in developing seeds. J. Exp. Bot. 52, 551–564. DOI: 10.1093/jexbot/52.356.551.

Olsen, O.-A., 2020: The modular control of cereal endosperm development. Trends Plant Sci., 25, 279–290. DOI: 10.1016/j.tplants.2019.12.003.

Pena, R.J., Nagarajan, P., Bates, L.S., 1982: Grain shrivelling in secondary hexaploid triticale. II. Morphology of mature and developing grains related to grain shrivelling. Cereal Chem. 59, 459–468.

Sabelli, P.A., Larkins, B.A., 2009: The development of endosperm in grasses. Plant Physiol. 149, 14–26. DOI: 10.1104/pp.108.129437.

Shewry, P.R., Mitchell, R.A.C., Tosi, P., Wan, Y., Underwood, C., Lovegrove, A., Freeman, J., Toole, G.A., Mills, E.N.C., Ward, J.L., 2012: An integrated study of grain development of wheat (cv. Hereward). J. Cereal Sci. 56, 21–30. DOI: 10.1016/j.jcs.2011.11.007.

Simmonds, D.H., O’Brien, T.P., 1981: Morphological and biochemical development of the wheat endosperm. Adv. Cereal Sci. Tech. 4, 5–70.

Sinha, N., Lynch, M., 1998: Fused organs in the adherent1 mutation in maize show altered epidermal walls with no perturbations in tissue identities. Planta 206, 184–195. DOI: 10.1007/s004250050390.

Stoddard, F.L., 1999: Survey of starch particle-size distribution in wheat and related species. Cereal Chem. 76, 145–149.

Tomaszewska, P., Kosina, R., 2018: Instability of endosperm development in amphiploids and their parental species in the genus Avena L. Plant Cell Rep. 37, 1145–1158. DOI: 10.1007/s00299018-2301-x.

Tomaszewska, P., Kosina, R., 2024: Oat species and interspecific amphiploids show predominance of diploid nuclei in the syncytial endosperm. J. Appl. Genet. 65, 1–11. DOI: 10.1007/s13353-023-00798-0.

Tonosaki, K., Ono, A., Kunisada, M., Nishino, M., Nagata, H., Sakamoto, S., Kijima, S.T., Furuumi, H., Nonomura, K.-I., Sato, Y., Ohme-Takagi, M., Endo, M., Comai, L., Hatakeyama, K., Kawakatsu, T., Kinoshita, T., 2021: Mutation of the imprinted gene OsEMF2a induces autonomous endosperm development and delayed cellularization in rice. Plant Cell 33, 85–103.

Vijayaraghavan, M.R., Prabhakar, K., 1984: The endosperm. In: Johri BM (ed) Embryology of Angiosperms. Springer, Berlin, pp 319–376.

Wang, Z., Li, W., Qi, J., Shi, P., Yin, Y., 2014: Starch accumulation, activities of key enzyme and gene expression in starch synthesis of wheat endosperm with different starch contents. J. Food Sci. Technol. 51, 419–429.

